# Contributions of white matter connectivity and BOLD modulation to cognitive aging: A lifespan structure-function association study

**DOI:** 10.1101/620443

**Authors:** Christina E. Webb, Karen M. Rodrigue, David A. Hoagey, Chris M. Foster, Kristen M. Kennedy

**Author notes:** Address correspondence to: K. M. Kennedy, 1600 Viceroy Dr., Suite 800, Dallas, TX, 75235, USA.

## Abstract

The ability to flexibly modulate brain activation to increasing cognitive challenge decreases with aging. This age-related decrease in dynamic range of function of regional gray matter may be, in part, due to age-related degradation of regional white matter tracts. Here, a lifespan sample of 171 healthy adults (aged 20-94) underwent MRI scanning including diffusion-weighted imaging (for tractography) and functional imaging (a digit *n*-back task). We utilized structural equation modeling to test the hypothesis that age-related decrements in white matter microstructure are associated with altered BOLD modulation, and both in turn, are associated with scanner-task accuracy and executive function performance. Specified structural equation model evidenced good fit, demonstrating that increased age negatively affects *n*-back task accuracy and executive function performance in part due to both degraded white matter tract microstructure and reduced task-difficulty related BOLD modulation. We further demonstrated that poorer white matter microstructure integrity was associated with weakened BOLD modulation, particularly in regions showing positive modulation effects, as opposed to negative modulation effects. This structure-function association study provides further evidence that structural connectivity influences functional activation, and the two mechanisms in tandem are predictive of cognitive performance, both during the task, and for cognition measured outside the scanner environment.

---

Dynamic range of blood-oxygen-level-dependent (BOLD) activity in response to parametrically increasing task demands decreases with age (Hakun and Johnson 2017; Kennedy et al. 2017; Rieck et al. 2017). Evidence from age comparison studies suggests that at low levels of task demands (or cognitive load) older adults increase their recruitment of cortical association regions in the prefrontal and parietal cortices, perhaps as a form of compensation to maintain similar levels of task performance as younger adults (Cappell et al. 2010; Schneider-Garces et al. 2010; Nagel et al. 2011; Garrett et al. 2013). While younger adults are able to increase activity in these regions to meet greater task demands, older adults do not maintain this increased recruitment, shifting to under-recruitment of prefrontal regions. Studies utilizing lifespan samples demonstrate that aging affects the ability to both positively modulate (i.e., increase activity) and negatively modulate (i.e., deactivate) in response to parametrically increasing cognitive demands, and that age-related reductions in dynamic range of modulation are associated with poorer cognitive performance (Kennedy et al. 2017; Rieck et al. 2017). In these studies age-related decreases in positive modulation are typically evident in task-related fronto-parietal regions (Kennedy et al. 2015, 2017; Rieck et al. 2017; see also Hakun and Johnson 2017, in an older adult sample), and are interpreted as reflecting reduced ability to successfully engage in strategic cognitive control processes under increasing task demands. These studies also demonstrate that aging is accompanied by decreased negative modulation to task difficulty (i.e., less deactivation with age) in regions typically associated with the default mode network (i.e., medial prefrontal cortex, anterior and posterior cingulate, precuneus, angular gyrus, and lateral temporal cortices; Persson et al. 2007; Park et al. 2010; Kennedy et al. 2017; Rieck et al. 2017), along with decreased functional connectivity among these regions (e.g., Sambataro et al. 2010). Failure to suppress default regions when task difficulty is high is suggested to reflect age-related dysregulation of resource allocation to cognitively-demanding tasks. Both up-modulation of fronto-parietal cognitive control regions and down-modulation of default network regions in response to cognitive challenge co-occur, suggesting synergistic relationships between the modulatory processes, especially in aging populations (Turner and Spreng 2015; Kennedy et al. 2017; Rieck et al. 2017). Overall, these findings suggest that individual differences in the capacity to flexibly modulate neural activity in response to increasing cognitive demands is a predictor of cognitive success, and that age-related decreases in modulation have negative cognitive consequences.

In addition to functional declines, aging is also accompanied by degradation of structural components of both gray and white matter. Gray matter volume and thickness, typically in heteromodal association cortices and hippocampus, decrease with age, and this age-related tissue loss has been associated with poorer cognitive functioning (Salat 2004; Raz and Kennedy 2009; Fjell and Walhovd 2010). In addition to alterations in volume and thickness of gray matter, white matter connections are particularly vulnerable to age-related degradation. Diffusion weighted imaging allows for estimation of the white matter fiber connectivity through measurement of microstructural properties of water diffusion. Fractional anisotropy (FA), an estimate of the degree of restricted diffusion of water molecules in white matter tracts, tends to show a negative association with age. This age-related decline is most notably observed within association tract fibers, which connect heteromodal gray matter regions, and may follow an anterior-to-posterior gradient in aging (Sullivan and Pfefferbaum 2006; Madden et al. 2009; Salat 2011). Age-related decreases in FA are indicative of declines in microstructural organization and conformation, which likely influence the efficiency of communication between gray matter regions connected by white matter pathways. In line with theories of cortical disconnection contributing to cognitive decline in aging (O’Sullivan et al. 2001; Bartzokis 2004), age-related decreases in FA have been associated with poorer cognitive performance, usually on tasks measuring aspects of executive processes (for review see Madden, et al., 2012). Furthermore, white matter integrity indices mediate relationships between age and cognitive performance (Kennedy and Raz 2009; Madden et al. 2009; Gold et al. 2010; Brickman et al. 2012; Samanez-Larkin et al. 2012; Borghesani et al. 2013), providing evidence that white matter connectivity alterations contribute to age-related decline in complex cognitive functions.

Communication among brain networks in support of successful cognitive performance is dependent, in part, upon intact white matter. Thus, age-related degradation of white matter tracts would be expected to impede functioning of cortical activity in brain regions connected by those pathways. Specifically, the dynamic range of regional gray matter function should be influenced by age-related alterations of white matter connectivity, and together, these neural properties should contribute to poorer cognitive functioning. However, relatively few studies have investigated the effects of age on the association between functional activity and white matter microstructural integrity. Existing studies provide evidence that individual differences in white matter connectivity are related to dysregulation of functional activation in middle-aged and older adults (Persson et al. 2006; Madden et al. 2007; de Chastelaine et al. 2011; Bennett and Rypma 2013; Daselaar et al. 2015; Hakun et al. 2015; Zhu et al. 2015; Brown et al. 2018). However, the directionality of these structure-function associations is somewhat mixed across studies and appears to be partially dependent upon the particular task and contrast used to assess functional activity. Additionally, white matter connectivity indices and functional activation tend to be localized from a few specific tracts and clusters, making it difficult to determine if these relationships are truly focal or whether they would also generalize across structural and functional ROIs.

It is also unclear how the aging process influences associations between white matter connectivity and functional activation, as most existing studies focus exclusively on older adults and/or do not directly compare these relationships between age groups. To date, only one study has examined structure-function relationships using DTI and fMRI simultaneously in a large continuous lifespan sample (Brown et al. 2015). In this study, age-related dysregulation of functional activation across the default mode network (DMN) was partially accounted for by reductions in FA of white matter connecting DMN regions. Importantly, this was only the case for the more difficult condition, and FA was not associated with DMN activation in the easier condition. A similar mediation was reported by Brown and colleagues (Brown et al. 2018) using separate younger and older age groups. Thus, alterations in white matter connectivity appear to be a plausible mechanism underlying age-related impairments in functional modulation to cognitive difficulty. It remains to be established whether parametric modulation of BOLD response to task demands is also related to white matter microstructure and whether these structure-function associations influence age-cognition associations across the lifespan.

Moreover, it is important to establish whether structure-function associations can explain age differences in cognitive function. Previous multimodal imaging studies have typically considered relationships between FA and behavior and BOLD activation and behavior separately, with age-related declines in FA associated with poorer performance and differences in BOLD activity differentially related to either better or poorer performance, depending on the task (Persson et al. 2006; Madden et al. 2007; Daselaar et al. 2015; Hakun et al. 2015; Zhu et al. 2015). Few studies have considered structure-function-behavior relationships simultaneously. de Chastelaine and colleagues (de Chastelaine et al. 2011) simultaneously included FA (from the genu of the corpus callosum) and BOLD response (from a cluster in right prefrontal cortex) in a linear regression model predicting task episodic memory performance in a group of older adults and found that both FA and functional activity accounted for unique variance in memory. Additionally, Brown and colleagues (Brown et al. 2015) noted unique effects of age, FA, and DMN activity on task accuracy and reaction time, but no interaction between FA and functional activation on performance. While cortical disconnection theories explaining age-related neural and cognitive decline imply that weakened structural connectivity contributes to dysregulation of brain function and impaired cognition, a complete test of this theory has yet to be established using multivariate methods.

Thus, the current study utilized structural equation modeling (SEM) to test the hypothesis that age-related structural connectivity degradation contributes to inefficient functional responses to increased task demands, which in turn affects cognitive performance. In an adult lifespan sample we measured 1) white matter tract microstructure across multiple tracts using tractography, 2) positive and negative BOLD modulation to difficulty associated with parametrically increasing working memory load on an *n*-back task using fMRI, and 3) cognitive performance both on *n*-back and executive function tasks. We included both brain regions evidencing positive modulation or negative modulation in our model to clarify whether individual differences in structural connectivity are specifically related to upregulation or downregulation of brain resources in response to task difficulty, as previous studies have shown mixed associations. The benefit of a structural equation modeling approach is that covariance among all variables are considered simultaneously, and thus hypothesized patterns of structure-function-cognition relationships can be tested concurrently.

## Materials and Methods

### Participants

Participants included 171 healthy adults (100 women), aged 20-94 years (*M*_age_ = 53.02, SD = 19.13) residing in the Dallas/Fort Worth metropolitan area. This sample was included in our previous report detailing age effects on dynamic range of BOLD modulation to cognitive difficulty (Kennedy et al. 2017). Participants were right-handed, native English speakers, had a minimum of high school education or equivalent (*M* = 15.59, SD = 2.49), and had normal or corrected-to-normal vision (when necessary vision was corrected to normal using MRI-compatible lenses). Participants were also free from a history of any metabolic, neurological or psychiatric disorders, head injury with loss of consciousness greater than 5 minutes, substance abuse, or cardiovascular disease (except for controlled hypertension), and did not have any contraindications for MRI. Screening was conducted to exclude individuals who might have either depression or dementia using the Center for Epidemiological Studies Depression Scale (CES-D ≥ 16; Radloff, 1977), and the Mini-Mental State Exam (MMSE < 26; Folstein, Folstein, & McHugh, 1975). Demographic characteristics divided by arbitrarily selected age groups are reported in Table 1; however, age was sampled and analyzed as a continuous variable in all statistical analyses. Participants provided written informed consent prior to study entry and all protocols were approved by the University of Texas at Dallas and the University of Texas Southwestern Medical Center institutional review boards.

**Table 1.**
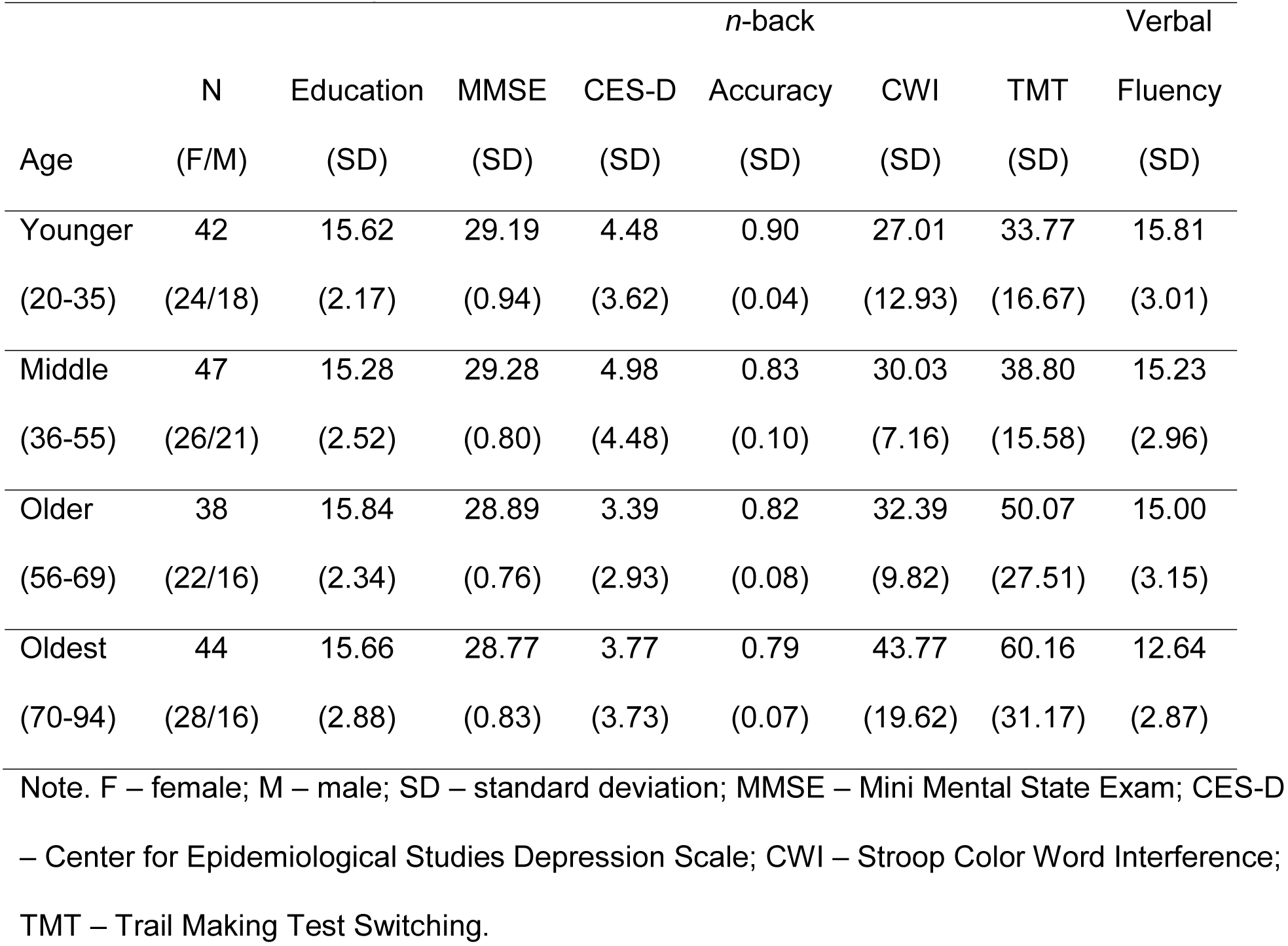
Participant Demographics and Task Performance

### MRI Data Acquisition

All participants were scanned on the same 3T Philips Achieva whole-body scanner equipped with a 32-channel head coil using SENSE encoding (Philips Medical Systems, Best, Netherlands). Sequences were completed during the same scanning session, which for this study included T1-weighted, diffusion-weighted, and task-based functional EPI sequences. The high-resolution anatomical images were acquired with a T1-weighted MP-RAGE sequence with the following parameters: 160 sagittal slices, 1 mm^3^ voxel size, FOV = 256 mm × 204 mm × 160 mm, TE = 3.8 ms, TR = 8.3 ms, flip angle = 12°, total time = 3:57 min. A diffusion tensor imaging (DTI) single shot EPI sequence was acquired with the following parameters: 65 axial slices with voxel size of 2 × 2 × 2.2 mm^3^ (reconstructed to 0.85 × 0.85 × 2.2 mm^3^), 30 diffusion-weighted directions (b-value = 1000s/mm^2^) with 1 non-diffusion weighted b_0_ (0 s/ mm^2^), TR/TE = 5608 ms/51 ms, FOV = 224 × 224, matrix = 112 × 112, total time = 4:19 min. Blood-oxygenation-level-dependent (BOLD) data were acquired with a T2*-weighted EPI sequence with 29 interleaved axial slices parallel to the AC-PC line, using the following parameters: 64 × 64 matrix, 3.4 × 3.4 × 5 mm^3^, FOV = 220 mm × 145 mm × 220 mm, TE = 30 ms, TR = 1500 ms, total time = 6:43 min per run × 3 runs. fMRI Task Procedure (*n*-back)

The scanner task consisted of a digit *n*-back working memory task with 4 levels of difficulty (0-back, 2-back, 3-back, 4-back). The task was presented in a blocked design consisting of 3 runs with 8 blocks each, including 2 blocks of each difficulty level. During each block, participants were shown a series of digits and were required to indicate whether the currently presented digit was the same or different as the one presented *n*-trials ago using an MRI-compatible button box (index finger – SAME; middle finger – DIFFERENT). At the beginning of each block participants were presented with a 5-second cue indicating the *n*-back condition for that block (0-back, 2-back, 3-back, 4-back), followed by a 2 second fixation, and then presentation of the series of digits. Digits (“2-9”) were presented for 500 ms, followed by a 2000 ms ISI in a pseudorandom order using Psychopy v1.77.02 (Peirce 2008). Refer to Kennedy et al. (2017) for a more detailed description of the *n*-back task procedure.

### fMRI Data Processing and Analysis

All preprocessing and statistical analyses were completed using SPM8 (Wellcome Department of Cognitive Neurology, London, UK) and in-house Matlab 2012b (Mathworks) scripts. ArtRepair toolbox (Mazaika et al. 2007) was used to identify outliers in EPI volumes due to motion (> 2mm motion displacement) or intensity shift (>3% deviation from the mean in global intensity spikes). In each participant, runs where >15% of volumes (∼40 volumes) were marked as outliers were removed from analyses, and participants were required to retain at least 2 (out of the 3) runs for inclusion. Data from 6 participants were excluded for the following reasons: excessive motion as identified via ArtRepair (*n* = 3), poor T1 quality (*n* = 2), >15% of no response trials (*n* = 1), MMSE < 26 (*n* = 1), leaving a final total of 171 participants, identical to the sample reported in a previous study from our lab (Kennedy et al. 2017). fMRI data were first corrected for differences in slice acquisition time and for within-run participant movement. Images were then normalized to standard stereotaxic MNI space, and smoothed with an 8mm isotropic FWHM Gaussian kernel.

fMRI data were analyzed in the general linear modeling (GLM) framework in SPM8. Individual subject responses to each level of difficulty (0-back, 2-back, 3-back, 4-back) were convolved with a canonical hemodynamic response function. ArtRepair estimates of motion for each subject were also included as nuisance regressors in the model. To measure modulation of BOLD activity in response to task difficulty, linear contrasts were created at the first-level (0-back < 2-back < 3-back < 4-back, using contrast weights of −2.25, −0.25, 0.75, 1.75). These contrast weightings were derived by subtracting the mean of the four *n*-back levels (2.25) from each level and were chosen to reflect the presumed step-wise increase in difficult of the *n*-back task. At the second-level, a voxel-wise linear regression was conducted with age as a continuous between-subjects covariate and the linear effect of task difficulty from the first-level as a within-subject variable predicting BOLD activity. We then examined both positive and negative effects of working memory load (i.e., areas that increase and decrease linearly with increasing working memory load, respectively). Second-level analyses were corrected at whole-brain FWE *p* < .05.

#### ROI Definition

Mean beta values for all significant clusters associated with positive modulation and negative modulation responses to parametric increases in task difficulty were extracted using MarsBaR (Brett et al. 2002). Clusters showing an effect of positive modulation (i.e., increasing activity with increasing working memory load), included bilateral dorsolateral prefrontal, parietal, and primary visual cortices, and bilateral cerebellum. In addition, clusters showing an effect of negative modulation (i.e., greater deactivation with increasing working memory load), included medial prefrontal cortex, bilateral temporal and occipital cortices, left parietal cortex, precuneus/posterior cingulate, and motor cortex (see Figure 1A). These regions of interest (ROIs) were used as observed indicators to construct two latent BOLD modulation factors (mean positive modulation, mean negative modulation) for use in the structural equation models. As spatial specificity was not the goal of the analysis, where appropriate, clusters in contralateral hemispheres and clusters that were in close spatial proximity were averaged to obtain a weighted mean based on cluster size of each region.

**Figure 1.**
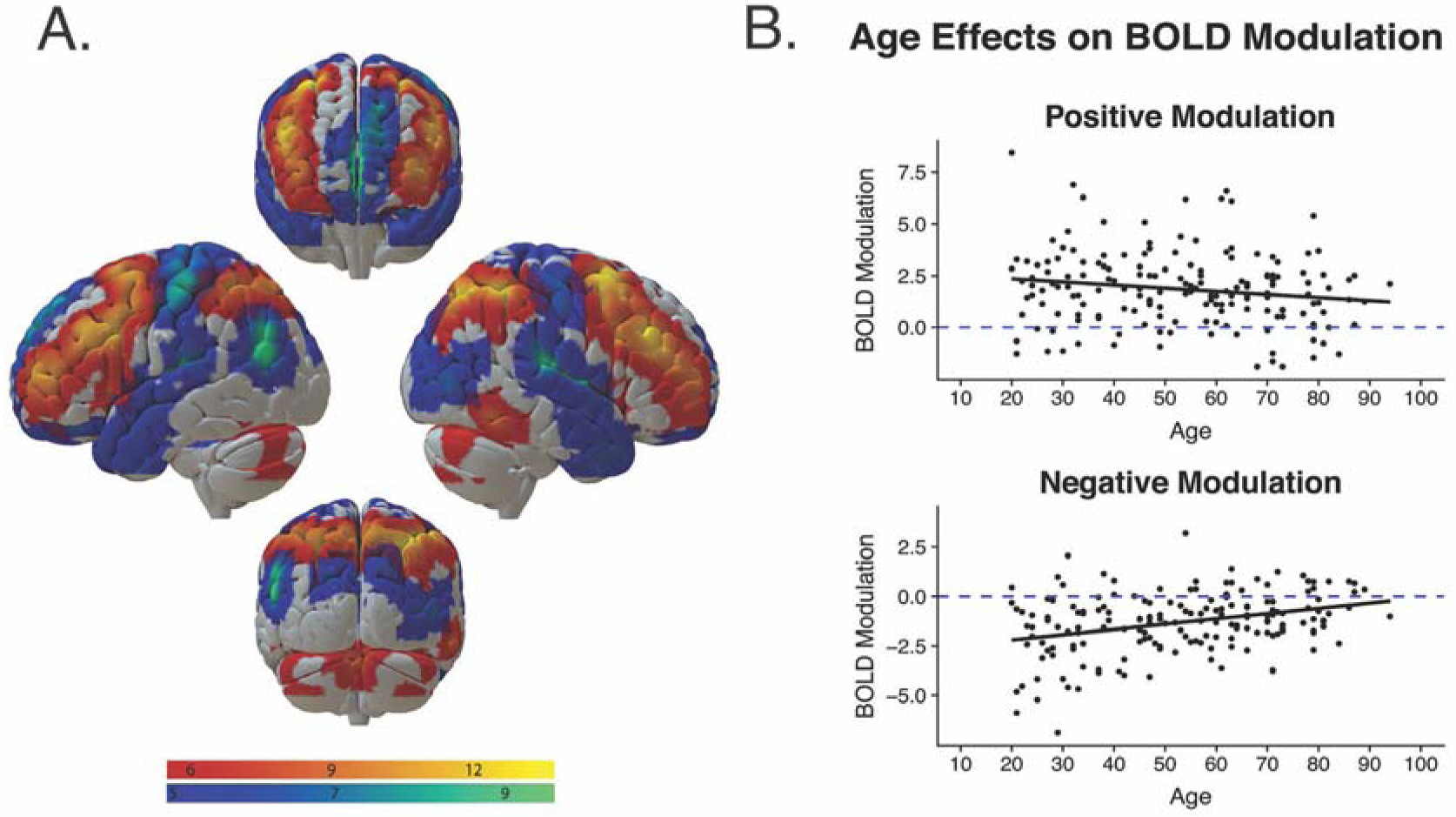
Effects of age on blood-oxygen-level-dependent (BOLD) modulation to difficulty. (A) BOLD activity modulation in response to increasing working memory load. Warm regions represent areas that evidenced increased activation in response to increasing working memory load (positive modulation) and cool regions represent areas that evidenced decreased activation in response to increasing working memory load (negative modulation). Clockwise from top: anterior view, right view, posterior view, left view. Whole-brain voxel threshold FWE *p* < .05. (B) Effects of age on positive (top) and negative (bottom) modulation of BOLD activation to working memory load, averaged across ROIs.

### DTI Data Processing and Analysis

Diffusion imaging data were preprocessed using the DTIPrep v1.2.4 quality control software suite to detect acquisition artifacts including susceptibility, eddy current, and subject movement distortions (Oguz et al. 2014). Using the default settings, slice-wise and gradient-wise artifacts, appearing as intensity distortions, were corrected by removing associated gradients from analysis. Appropriate corrections were applied to minimize the effects of distortions, including those caused by head motion in the scanner, by removing gradients determined to be of insufficient quality, at the default threshold levels, and by registering all remaining gradients to the non-weighted b_0_ image. On average, only four gradients were removed per subject. Diffusion directions were adjusted to account for independent rotations of any gradient relative to the original encoding direction (Leemans and Jones 2009). Diffusion tensors were calculated using the DSI Studio software package build from September 26^th^, 2014 (Yeh et al. 2013; http://dsi-studio.labsolver.org). Deterministic tractography, implemented in DSI Studio, was utilized to construct white matter tracts of interest that best represented association tract bundles, as well as representing patterns of connectivity covering regions supporting working memory and general higher-order cognition. These six tracts of interest consisted of the cingulum bundle, bilateral superior longitudinal fasciculus (SLF), inferior fronto-occipital fasciculus (IFOF), inferior longitudinal fasciculus (ILF), uncinate fasciculus (UF), and the genu of the corpus collosum (Genu). Fractional anisotropy (FA) was averaged across all voxels in each tract and extracted as the diffusion metric of interest. Tractography was restricted to voxels containing FA values > .20 for all participants. A deterministic tracking algorithm implemented in DSI Studio (Yeh et al. 2013) was used with the following parameters: maximum turning angle of 60°, step size of 1 mm, and a minimum/maximum length of 20/500 mm, respectively. All tract output was visually inspected and those with streamline counts 1.4 standard deviations below the mean were excluded from analysis. Deterministic tractography of the bilateral tracts was conducted by creating anatomical seeds and regions of inclusion and avoidance, based on consultation with atlases (Desikan et al. 2006; Hua et al. 2008; Mori et al. 2008), on the 1mm MNI template and warping each region into subject diffusion space as specified below for each tract.

#### Cingulum

Tracking of the cingulum was accomplished using the cingulate gyrus from the JHU atlas as the only ROI. Regions of avoidance were necessary as follows: to remove fibers tracking medially through the corpus collosum with a midsagittal plane, or laterally with parasagittal slices on either side of the ROIs; to remove fibers tracking into subcortical regions with planes encompassing all coronal slices directly below the corpus callosum, and to remove aberrant crossing fibers from the centrum semiovale with a coronal plane behind the centrum semiovale.

#### SLF

Tracking of the SLF was accomplished using two coronal inclusionary slices placed in the white matter of the parietal lobe which, when viewed sagittally, would align with the anterior and posterior ends of the splenium of the corpus callosum (*y* = −27 and *y* = −46). Additionally, to restrict our analyses to the fronto-parietal portion of the SLF, an axial exclusionary plane was created below the parietal white matter to remove any fibers extending into the arcuate fasciculus (*z* = 14). Finally, to remove commissural and projection fibers, a sagittal exclusionary plane was drawn medial to the parietal white matter (*x* = 20).

#### IFOF

Tracking of the IFOF was accomplished using whole plane coronal inclusionary slices that ensured each tract extended between the frontal (posterior to the genu of the corpus callosum) and the occipital lobes (posterior to the posterior thalamic radiation). To restrict spurious fibers from analyses, axial exclusionary slices were added above the corpus callosum (*z* = 35) and below the frontal and occipital lobes (*z* = −24). Additional regions of exclusion were added to remove fibers extending into the internal and external capsules.

#### ILF

Tracking of the ILF was accomplished using coronal regions of inclusion placed just posterior to the temporal pole and in the occipital lobe. To remove portions of fibers which extended into the frontal lobe, two exclusionary regions were placed: one coronal plane (posterior to the genu of the corpus callosum) and one sagittal plane (placed medial to the temporal pole and only extended to the anterior).

#### UF

Tracking of the UF was accomplished using two cubic regions of inclusion (approximately 15 mm^3^); one placed in the superior portion of the temporal pole and the other placed posterior to the frontal pole in the white matter medial to the orbitofrontal cortex. Additionally, two exclusionary planes were drawn to prevent spurious fibers: one coronal plane at *y* = −15 to prevent fibers tacking into the ILF, and one axial plane at *z* = 25 to prevent fibers from turning in the superior direction.

#### Genu

Tracking of the corpus callosum genu was accomplished using a hand-drawn genu ROI, based off of the vertical subdivision scheme proposed by Hofer and Frahm (Hofer and Frahm 2006) and the anatomically based classification system devised by Witelson (Witelson 1989). Additionally, this was supplemented by a midsagittal ROI to ensure that all fibers were traversing into each hemisphere. Three regions of avoidance were used: one coronal plane placed posterior to the genu to prevent fibers from tracking into other segments of the corpus callosum; parasagittal slices at the edges of the ROI to prevent spurious fibers extending laterally, and the cingulum ROIs to remove any spurious fibers from the cingulum.

### Cognitive Performance

#### n-back Task

Response time (RT) and accuracy data were recorded for each trial. Average accuracy across all trials was calculated for each level of working memory load in the *n*-back fMRI task (0-, 2-, 3-, 4-back) for each participant. Mean accuracy scores (percent correct on all trials) for each of the four load levels were used as observed indicators to create a latent construct representing accuracy on the *n*-back task.

#### Executive Function Tests

Prior to the scanning visit, participants completed two sessions consisting of a wide battery of cognitive and psychometric testing. To test whether brain variables account for age-related variance in cognition measured independent of the fMRI task, several measures of switching and inhibition from the Delis-Kaplan Executive Function System (D-KEFS; Delis et al. 2001) were used as observed indicators of a latent Executive Function variable. These included the Color Word Interference/Stroop (CWI), Trail Making Test (TMT), and Verbal Fluency switching tasks. Performance on the Color Word Interference task was calculated as the time taken in seconds for switching and inhibition conditions, adjusted for baseline rates of color naming and word reading. Trails performance was measured as the time taken to complete number/letter switching, adjusted for performance on number and letter sequencing tasks alone. Verbal fluency was defined as total accuracy on the category switching task. Scores on the Color Word Interference task and Trail Making Test were multiplied by −1 to scale all scores in the same direction, such that higher scores reflect greater executive function.

### Statistical Analyses

#### Measurement Model Construction

Prior to specification of the structural models, latent variables were created to represent both Positive and Negative BOLD Modulation, FA, *n*-back Accuracy and Executive Function. These latent variables represent shared individual differences among the indicators (tracts, ROIs, or cognition) of each variable. As mentioned previously, mean beta values extracted from four bilateral ROIs showing an effect of positive modulation (i.e., increased BOLD activity as a function of increasing working memory load) served as indicators of a latent Positive Modulation factor. The same procedure was implemented for negative modulation, where mean beta values from six ROIs showing an effect of negative modulation (i.e., increased deactivation as a function of increasing working memory load) were used as indicators of a latent Negative Modulation variable. FA values from the six white matter tracts served as indicators of a latent variable representing FA. Accuracy on each of the four levels of difficulty during the *n*-back task were chosen as indicators of a latent variable representing *n*-back Accuracy, and scores on each of the three executive function tests served as indicators of a latent Executive Function variable.

#### Structural equation model specification

Structural equation modeling was used to simultaneously estimate relationships among age, white matter microstructure, BOLD modulation, and cognition (*n*-back accuracy and executive function). Structural equation modeling was conducted with Mplus software version 8 (Muthen and Muthen 2017) using maximum likelihood estimation. Maximum likelihood estimation allows for all participants to be included in the model, despite missing data on some of the variables (*n* = 1 missing DTI data). Goodness of fit was assessed using the following indices: Root Mean Square Error of Approximation (RMSEA) < 0.08, Comparative Fit Index (CFI) and Tucker-Lewis Index (TLI) > 0.90, and Standardized Root Mean Square Residual (SRMR) < 0.08 (MacCallum et al. 1996; Hu and Bentler 1999; Kline 2011). Significance of direct and indirect paths was determined based on 95% confidence intervals resulting from bootstrapping with 5,000 samples. Confidence intervals that did not contain zero were considered significant.

A structural equation model was tested to investigate the theory that age-related variance in cognition is influenced by global white matter degradation and functional modulation. Age served as the exogenous variable and latent variables characterizing both *n*-back Accuracy and an out-of-scanner measure of Executive Function were the primary cognitive outcome variables. Latent variables representing FA, Positive, and Negative BOLD Modulation served as mediating variables. As age is consistently shown to relate to FA and BOLD modulation, as well as cognition, paths from age to all latent factors were estimated. In accord with theory suggesting that age-related decline in brain structure affects gray matter function, a path from age to both Positive and Negative Modulation was estimated with FA as the mediating variable. Positive and Negative Modulation were allowed to covary as previous research indicates the coupling of positive and negative modulation to difficulty (Kennedy et al. 2017; Rieck et al. 2017). Lastly, paths were estimated from FA, Positive Modulation, and Negative Modulation to both *n*-back Accuracy and Executive Function. All factor means were fixed to zero and factor variances were fixed to one to standardize the factors. All factor loadings were allowed to be freely estimated.

## Results

### Age Effects on Brain Function, Structure, and Cognition

#### Age and BOLD Modulation

Table 2 depicts the observed zero-order Pearson correlations among all variables included in the model. Age was generally associated with reduced activity in regions showing a positive modulation response to task difficulty (warm scale in Figure 1A), but only significantly in the lateral prefrontal and superior parietal ROIs. Age was also significantly associated with increased activity in all regions showing a negative modulation response to task difficulty (cool scale in Figure 1A), indicating reduced deactivation in these regions with increasing age. Figure 1B depicts age trends averaged across ROIs showing effects of positive and negative modulation.

**Table 2.** Bivariate correlations among observed variables

|  | 1 | 2 | 3 | 4 | 5 | 6 | 7 | 8 | 9 | 10 | 11 | 12 | 13 | 14 | 15 | 16 | 17 | 18 | 19 | 20 | 21 | 22 |
| --- | --- | --- | --- | --- | --- | --- | --- | --- | --- | --- | --- | --- | --- | --- | --- | --- | --- | --- | --- | --- | --- | --- |
| 1. Age |  |  |  |  |  |  |  |  |  |  |  |  |  |  |  |  |  |  |  |  |  |  |
| 2. n0 | **-0.17** |  |  |  |  |  |  |  |  |  |  |  |  |  |  |  |  |  |  |  |  |  |
| 3. n2 | **-0.44** | **0.44** |  |  |  |  |  |  |  |  |  |  |  |  |  |  |  |  |  |  |  |  |
| 4. n3 | **-0.53** | **0.35** | **0.82** |  |  |  |  |  |  |  |  |  |  |  |  |  |  |  |  |  |  |  |
| 5. n4 | **-0.46** | **0.40** | **0.74** | **0.80** |  |  |  |  |  |  |  |  |  |  |  |  |  |  |  |  |  |  |
| 6. CWI | **-0.44** | 0.12 | **0.36** | **0.36** | **0.33** |  |  |  |  |  |  |  |  |  |  |  |  |  |  |  |  |  |
| 7. TMT | **-0.38** | 0.07 | **0.50** | **0.44** | **0.39** | **0.43** |  |  |  |  |  |  |  |  |  |  |  |  |  |  |  |  |
| 8. Verbal | **-0.37** | 0.07 | **0.35** | **0.34** | **0.29** | **0.28** | **0.39** |  |  |  |  |  |  |  |  |  |  |  |  |  |  |  |
| 9. Cingulum | **-0.60** | **0.19** | **0.37** | **0.37** | **0.34** | **0.28** | **0.28** | **0.28** |  |  |  |  |  |  |  |  |  |  |  |  |  |  |
| 10. SLF | **-0.61** | **0.18** | **0.31** | **0.34** | **0.29** | **0.34** | **0.27** | **0.25** | **0.85** |  |  |  |  |  |  |  |  |  |  |  |  |  |
| 11. IFOF | **-0.56** | 0.12 | **0.33** | **0.33** | **0.26** | **0.36** | **0.33** | **0.27** | **0.76** | **0.76** |  |  |  |  |  |  |  |  |  |  |  |  |
| 12. ILF | **-0.59** | **0.16** | **0.33** | **0.36** | **0.29** | **0.37** | **0.36** | **0.27** | **0.78** | **0.81** | **0.87** |  |  |  |  |  |  |  |  |  |  |  |
| 13. UF | **-0.66** | 0.13 | **0.38** | **0.36** | **0.34** | **0.32** | **0.33** | **0.29** | **0.78** | **0.76** | **0.75** | **0.77** |  |  |  |  |  |  |  |  |  |  |
| 14. Genu | **-0.71** | 0.10 | **0.35** | **0.40** | **0.34** | **0.32** | **0.34** | **0.33** | **0.85** | **0.80** | **0.77** | **0.77** | **0.82** |  |  |  |  |  |  |  |  |  |
| 15. Pos. Frontal | **-0.18** | **0.22** | **0.32** | **0.29** | **0.26** | **0.16** | 0.13 | **0.17** | **0.25** | **0.25** | **0.20** | **0.25** | **0.21** | 0.20 |  |  |  |  |  |  |  |  |
| 16. Pos. Parietal | **-0.21** | **0.23** | **0.32** | **0.30** | **0.28** | **0.17** | **0.18** | **0.22** | **0.24** | **0.26** | **0.20** | **0.25** | **0.21** | **0.19** | **0.91** |  |  |  |  |  |  |  |
| 17. Pos. Occipital | -0.10 | -0.03 | 0.03 | 0.13 | 0.04 | 0.03 | 0.08 | 0.12 | 0.09 | 0.10 | **0.17** | 0.13 | 0.06 | 0.06 | **0.49** | **0.52** |  |  |  |  |  |  |
| 18. Pos. Cerebellum | -0.08 | **0.15** | **0.26** | **0.23** | **0.20** | **0.16** | **0.16** | **0.15** | 0.15 | **0.16** | 0.13 | 0.15 | 0.12 | 0.10 | **0.81** | **0.82** | **0.53** |  |  |  |  |  |
| 19. Neg. Frontal | **0.27** | **-0.17** | **-0.24** | **-0.26** | **-0.27** | -0.08 | **-0.22** | -0.15 | -0.15 | -0.13 | -0.07 | -0.10 | -0.12 | -0.14 | **0.31** | **0.20** | **0.21** | **0.33** |  |  |  |  |
| 20. Neg. Motor | **0.18** | -0.12 | **-0.17** | **-0.18** | -0.14 | -0.15 | **-0.25** | -0.14 | **-0.16** | -0.13 | -0.12 | -0.12 | -0.15 | **-0.15** | **0.18** | **0.19** | **0.27** | **0.27** | **0.36** |  |  |  |
| 21. Neg. Temporal | **0.25** | **-0.19** | **-0.26** | **-0.23** | **-0.25** | **-0.19** | **-0.32** | **-0.18** | **-0.17** | -0.12 | -0.14 | -0.13 | **-0.16** | **-0.19** | **0.34** | **0.32** | **0.43** | **0.40** | **0.55** | **0.65** |  |  |
| 22. Neg. Precuneus | **0.26** | **-0.17** | **-0.25** | **-0.19** | **-0.22** | **-0.16** | **-0.23** | -0.10 | **-0.18** | **-0.17** | -0.11 | **-0.16** | **-0.18** | **-0.21** | **0.45** | **0.45** | **0.56** | **0.47** | **0.58** | **0.59** | **0.80** |  |
| 23. Neg. Parietal | **0.33** | -0.11 | **-0.28** | **-0.22** | **-0.25** | **-0.24** | **-0.18** | -0.05 | **-0.33** | **-0.28** | **-0.29** | **-0.26** | **-0.29** | **-0.36** | **0.31** | **0.34** | **0.38** | **0.38** | **0.54** | **0.43** | **0.64** | **0.74** |
| 24. Neg. Occipital | **0.40** | **-0.18** | **-0.26** | **-0.26** | **-0.29** | **-0.17** | **-0.25** | -0.15 | **-0.24** | **-0.27** | **-0.21** | **-0.25** | **-0.25** | **-0.32** | **0.27** | **0.27** | **0.50** | **0.35** | **0.51** | **0.56** | **0.71** | **0.73** |

Note. Bold font indicates significant Pearson correlation at $p < .05$ ; $n0$ – 0-back accuracy; $n2$ – 2-back accuracy; $n3$ – 3-back accuracy; $n4$ – 4-back accuracy; CWI – Stroop Color Word Interference task; TMT – Trail Making Test; SLF – superior longitudinal fasciculus; IFOF -- inferior fronto-occipital fasciculus; ILF – inferior longitudinal fasciculus, UF – uncinate fasciculus, Genu – genu of the corpus callosum; Pos. – positive modulation; Neg. – negative modulation.

**Table 3.**
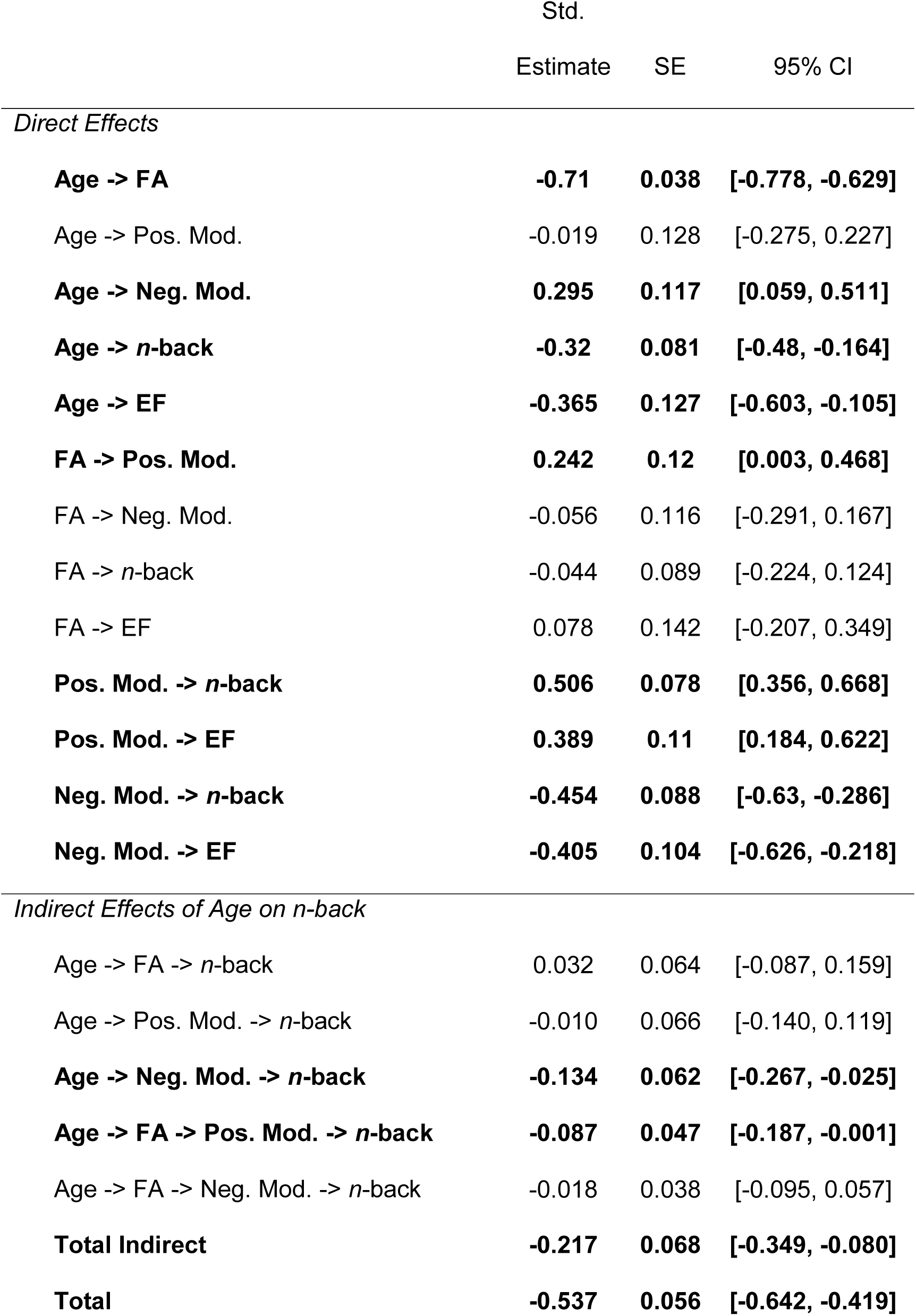
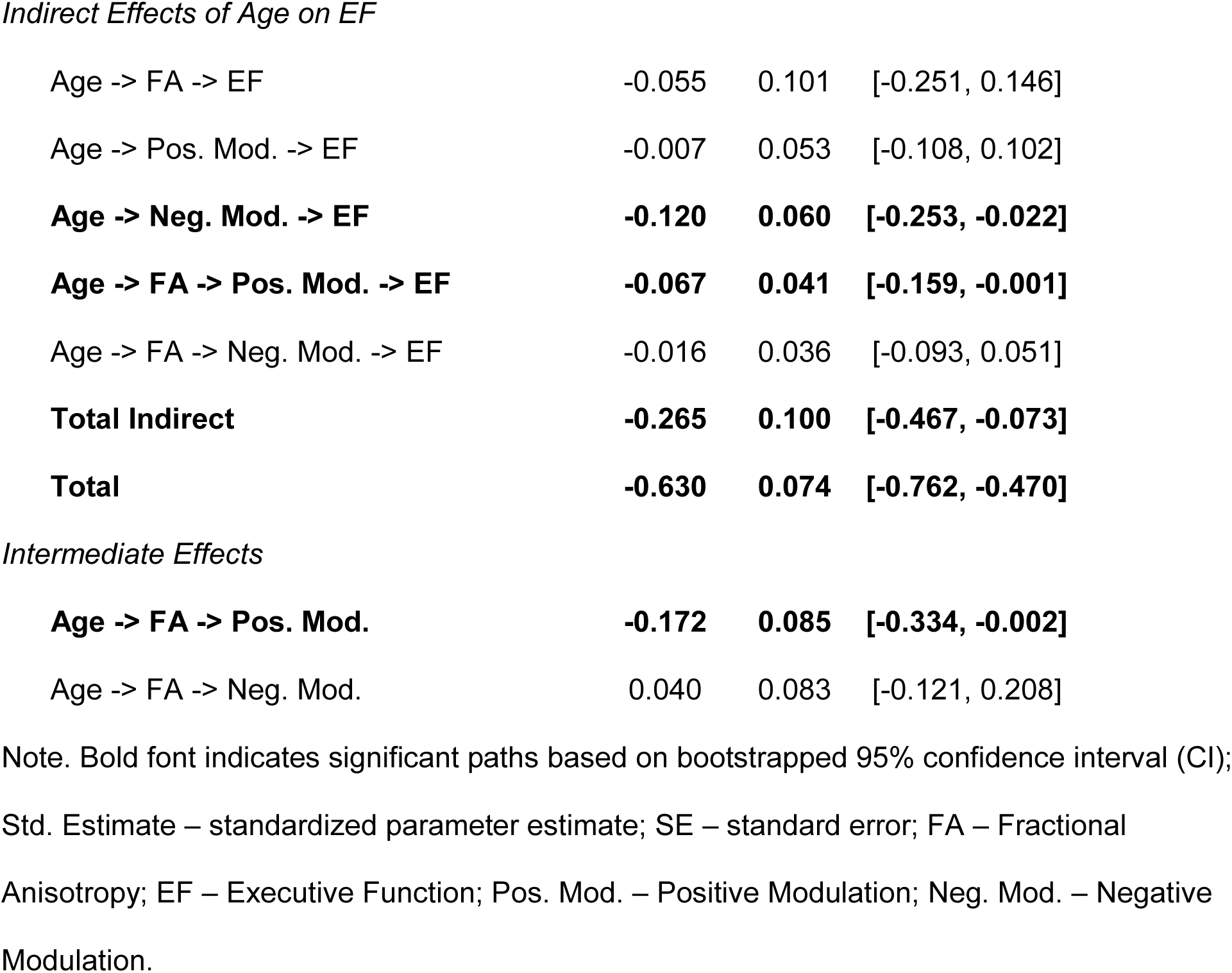
Effects of Age on n-back Accuracy and Executive Function

|  | Std.<br>Estimate | SE | 95% CI |
| --- | --- | --- | --- |
| *Direct Effects* |  |  |  |
| **Age -> FA** | **-0.71** | **0.038** | **[-0.778, -0.629]** |
| Age -> Pos. Mod. | -0.019 | 0.128 | [-0.275, 0.227] |
| **Age -> Neg. Mod.** | **0.295** | **0.117** | **[0.059, 0.511]** |
| **Age -> n-back** | **-0.32** | **0.081** | **[-0.48, -0.164]** |
| **Age -> EF** | **-0.365** | **0.127** | **[-0.603, -0.105]** |
| **FA -> Pos. Mod.** | **0.242** | **0.12** | **[0.003, 0.468]** |
| FA -> Neg. Mod. | -0.056 | 0.116 | [-0.291, 0.167] |
| FA -> n-back | -0.044 | 0.089 | [-0.224, 0.124] |
| FA -> EF | 0.078 | 0.142 | [-0.207, 0.349] |
| **Pos. Mod. -> n-back** | **0.506** | **0.078** | **[0.356, 0.668]** |
| **Pos. Mod. -> EF** | **0.389** | **0.11** | **[0.184, 0.622]** |
| **Neg. Mod. -> n-back** | **-0.454** | **0.088** | **[-0.63, -0.286]** |
| **Neg. Mod. -> EF** | **-0.405** | **0.104** | **[-0.626, -0.218]** |
| *Indirect Effects of Age on n-back* |  |  |  |
| Age -> FA -> n-back | 0.032 | 0.064 | [-0.087, 0.159] |
| Age -> Pos. Mod. -> n-back | -0.010 | 0.066 | [-0.140, 0.119] |
| **Age -> Neg. Mod. -> n-back** | **-0.134** | **0.062** | **[-0.267, -0.025]** |
| **Age -> FA -> Pos. Mod. -> n-back** | **-0.087** | **0.047** | **[-0.187, -0.001]** |
| Age -> FA -> Neg. Mod. -> n-back | -0.018 | 0.038 | [-0.095, 0.057] |
| **Total Indirect** | **-0.217** | **0.068** | **[-0.349, -0.080]** |
| **Total** | **-0.537** | **0.056** | **[-0.642, -0.419]** |
| *Indirect Effects of Age on EF* |  |  |  |
| Age -> FA -> EF | -0.055 | 0.101 | [-0.251, 0.146] |
| Age -> Pos. Mod. -> EF | -0.007 | 0.053 | [-0.108, 0.102] |
| **Age -> Neg. Mod. -> EF** | **-0.120** | **0.060** | **[-0.253, -0.022]** |
| **Age -> FA -> Pos. Mod. -> EF** | **-0.067** | **0.041** | **[-0.159, -0.001]** |
| Age -> FA -> Neg. Mod. -> EF | -0.016 | 0.036 | [-0.093, 0.051] |
| **Total Indirect** | **-0.265** | **0.100** | **[-0.467, -0.073]** |
| **Total** | **-0.630** | **0.074** | **[-0.762, -0.470]** |
| *Intermediate Effects* |  |  |  |
| **Age -> FA -> Pos. Mod.** | **-0.172** | **0.085** | **[-0.334, -0.002]** |
| Age -> FA -> Neg. Mod. | 0.040 | 0.083 | [-0.121, 0.208] |

Note. Bold font indicates significant paths based on bootstrapped 95% confidence interval (CI);
Std. Estimate – standardized parameter estimate; SE – standard error; FA – Fractional
Anisotropy; EF – Executive Function; Pos. Mod. – Positive Modulation; Neg. Mod. – Negative Modulation.

#### Age and White Matter Tract FA

The mean (± standard deviation) of FA values, averaged across hemisphere, for each tract in the subset of participants with DTI data (*n* = 170) were: cingulum = 0.39 (0.02), SLF = 0.40 (0.02), IFOF = 0.45 (0.02), ILF = 0.44 (0.02), UF = 0.39 (0.02), genu = 0.45 (0.03). Older age was associated with significantly reduced FA in all measured white matter tracts (*r*s ranged from −.56 to −.71 across the tracts; see Figure 2 and Table 2).

**Figure 2.**
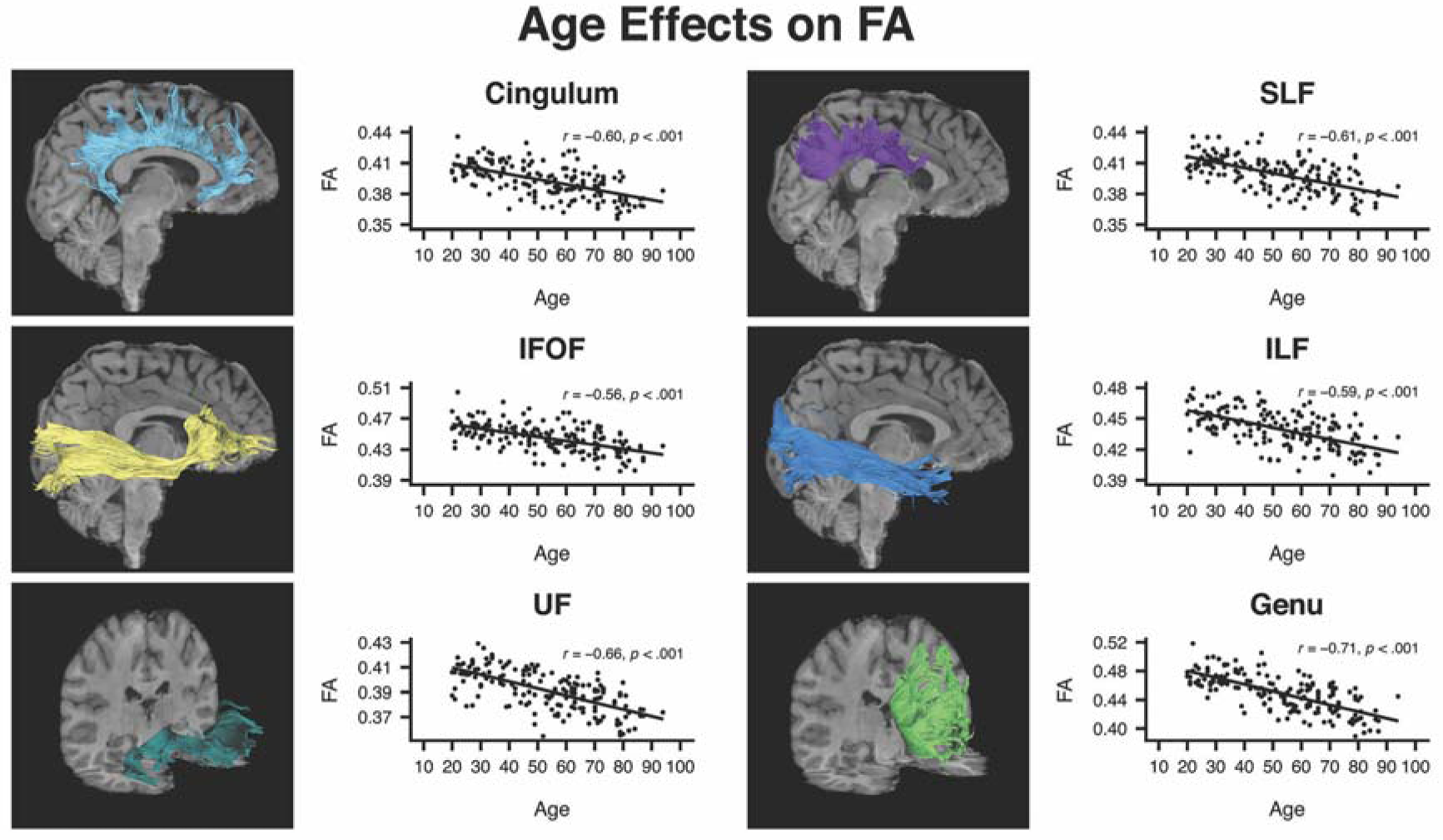
Sample white matter tract images from a representative participant and correlations between fractional anisotropy (FA) and age for cingulum, superior longitudinal fasciculus (SLF), inferior fronto-occipital fasciculus (IFOF), inferior longitudinal fasciculus (ILF), uncinate fasciculus (UF), genu of the corpus callosum (Genu).

#### Age and Cognitive Task Performance

Mean accuracy on the fMRI *n*-back task and mean performance on executive function tasks are reported in Table 1, broken down by arbitrary age group. Accuracy on all four levels of the fMRI *n*-back task decreased significantly with age (see Table 2). A repeated measures GLM with *n*-back accuracy as a within-subject factor with 4 levels (0-back, 2-back, 3-back, 4-back) and mean-centered age as a between-subjects factor revealed both a main effect of age [*F*(1,169) = 50.29, *p* < .001] and difficulty [*F*(3,507) = 280.08, *p* < .001]. Importantly, there was an age × working memory load interaction [*F*(3,507) = 12.72, *p* < .001], indicating that the negative effect of difficulty on accuracy became stronger with increasing age. Performance on all executive function tasks also decreased significantly with age, detailed in Table 2 (note that both Color Word Interference and Trail Making Test were reverse-scored; higher scores represent greater executive function).

#### Cognitive and Brain Variable Associations

Across all participants, accuracy on the *n*-back task was largely negatively correlated with regions showing negative modulation and positively correlated with regions showing positive modulation (Table 2), indicating that greater range of modulation to task difficulty was associated with higher accuracy on the *n*-back task. Accuracy on *n*-back was also positively correlated with FA values in all tracts, indicating that greater white matter microstructural integrity (as measured via FA) was associated with greater task accuracy. The same patterns of relationships were observed between performance on the executive function tasks and modulation, and between executive function and FA. Lastly, across participants, greater FA was associated with greater positive modulation and lower negative modulation in response to task difficulty, and both positive and negative modulation were highly correlated.

#### Structural Equation Models

The final SEM is depicted in Figure 3, with solid lines representing significant paths. Squares represent observed continuous variables and circles represent latent variables. The hypothesized model demonstrated good fit to the data [RMSEA: 0.067 [0.056, 0.077], CFI: 0.945, TLI: 0.936, SRMR: 0.055]. All standardized estimates, standard errors, and 95% confidence intervals for direct and indirect effects in the model are presented in Table 4. Age was negatively related to FA, *n*-back Accuracy, and Executive Function, and positively related to Negative Modulation. Age was not directly associated with Positive Modulation in this model. FA was significantly related to Positive (Std. Est. [95% CI] = 0.242 [0.003, 0.468]), but not Negative Modulation. Both Positive and Negative Modulation to difficulty were directly related to both *n-*back Accuracy (Std. Est. [95% CI]: Pos. Mod. = −.0.506 [0.356, 0.668]; Neg. Mod. = −0.454 [−0.62, −0.286]) and Executive Function (Std. Est. [95% CI]: Pos. Mod. = 0.389 [0.184, 0.622]; Neg. Mod. = −0.405 [−0.626, −0.218]); however no significant direct relationship between FA and either *n*-back Accuracy or Executive Function was identified. Tests of indirect effects indicated that the total indirect effect of age on both *n*-back Accuracy and Executive Function through white matter tract FA was significant (Std. Indirect Est. [95% CI]: *n*-back = −0.217 [−0.349, −0.08]; EF = −0.265 [−0.467, −0.073]), and that age exerted small specific indirect effects on *n*-back Accuracy and Executive Function through FA and Positive Modulation (Std. Indirect Est. [95% CI]: *n*-back = −0.087 [−0.187, −0.001]; EF = −0.067 [−0.159, −0.001]). In addition, age had a specific indirect effect on both Accuracy and Executive Function through its effect on Negative Modulation (Std. Indirect Est. [95% CI]: *n*-back = −0.134 [−0.267, −0.025]; EF = −0.12 [−0.253, −0.022]), such that greater age was associated with reduced FA, which was associated with poorer working memory and executive function. There was also a significant intermediate indirect effect of age on Positive Modulation through FA (Std. Indirect Est. [95% CI] = −0.172 [−0.334, −0.002]), but no indirect effect of age on Negative Modulation via FA. Overall, this model suggests that age-related degradation of white matter connections and reductions in dynamic range of brain activity, together, are associated with poorer cognitive performance, and highlights the critical role of white matter microstructure in supporting regulation of brain activation and cognitive function. Importantly, these relationships were also observed when extending to executive function tasks that were independent of functional activation.

**Figure 3.**
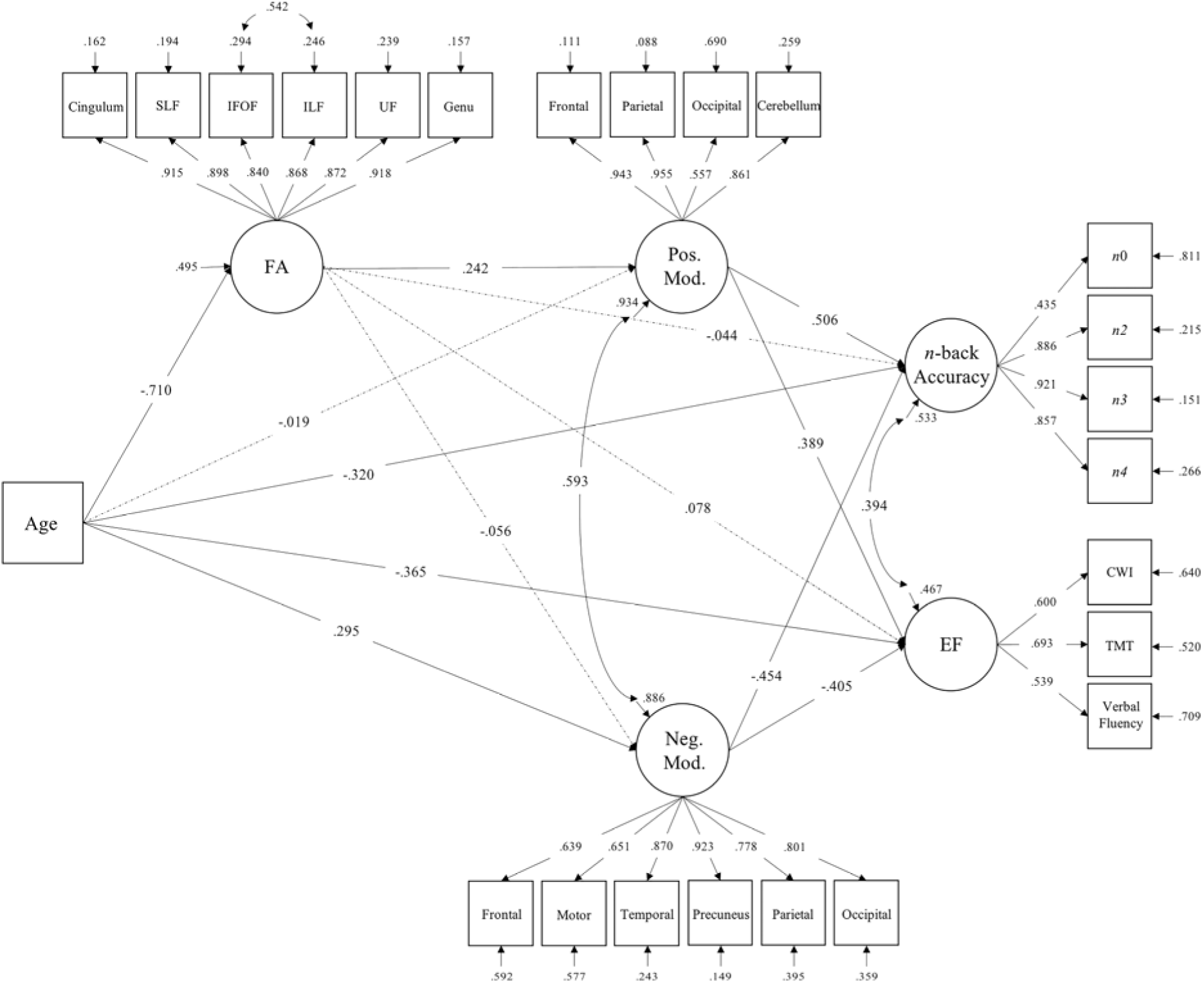
Structural Equation Model Predicting *n*-back Accuracy and Executive Function from Age, Tractography FA, and BOLD Modulation to Difficulty. Solid lines represent significant paths in the model at 95% confidence interval. Dashed lines represent non-significant estimated paths. Path values represent standardized parameter estimates. FA – Fractional Anisotropy; SLF – superior longitudinal fasciculus; IFOF – inferior fronto-occipital fasciculus; ILF – inferior longitudinal fasciculus; UF – uncinate fasciculus; Genu – genu of the corpus callosum; Pos. Mod. – Positive Modulation; Neg. Mod. – Negative Modulation; *n*0 – 0-back accuracy; *n*2 – 2-back accuracy; *n*3 – 3-back accuracy; *n*4 – 4-back accuracy; EF – Executive Function; CWI – Stroop Color Word Interference; TMT – Trail Making Test.

## Discussion

A wealth of previous studies have separately described structural or functional influences on age-related cognitive decline; yet there has been little work directly testing the assumption that structural disconnection accompanying aging contributes to functional dysregulation and exerts negative effects on cognition. The present study tested this hypothesis by simultaneously examining structure-function-cognition relationships using SEM in a lifespan sample of healthy adults. Our results provide evidence that age-related degradation of white matter connectivity specifically influences the magnitude of positive BOLD modulation to task difficulty, which contributes to age-related differences in cognitive performance. Additionally, we found that negative BOLD modulation independently explained age-related variation in cognition, such that greater age was associated with less modulation of deactivation and was associated with reduced cognitive performance.

Replicating previous research, we observed effects of both positive and negative BOLD activity modulation in response to greater working memory load across all participants. Effects of positive modulation, reflecting increased activity in response to greater cognitive load, were observed in regions typically thought to be part of the cognitive control network and often engaged in tasks of working memory (bilateral fronto-parietal, cerebellum, early visual cortex; Cabeza and Nyberg 2000; Owen et al. 2005; Nagel et al. 2011). In contrast, effects of negative modulation were observed in regions often associated with the default mode network (medial prefrontal, posterior cingulate/precuneus, bilateral lateral temporal cortices; Gusnard and Raichle 2001; Raichle et al. 2001; Persson et al. 2007; Park et al. 2010). Importantly, we observed age-related differences in this dynamic modulation of functional activity to cognitive challenge. Both the ability to increase activity in regions showing positive modulation effects and the ability to decrease activity in regions showing negative modulation effects decreased as a function of age. Weakened positive and negative functional modulation to cognitive demand in healthy aging has been documented across various tasks (Persson et al. 2007; Park 2010; Garrett et al. 2013; Hakun and Johnson 2017; Kennedy et al. 2017; Rieck et al. 2017), and is thought to reflect age-related reductions in the ability to flexibly engage neural resources to meet task demands. Specifically, reductions in positive modulation with advancing age imply that aging limits the ability to upregulate task-relevant control regions to successfully perform a task, while reductions in negative modulation with age reflect less suppression of default processes under greater cognitive load in aging (Persson et al. 2007; Sambataro et al. 2010).

In addition to reduced functional modulation range with age, we also observed age-associated declines in white matter fiber FA across all tracts of interest, suggesting decreased connectivity. Reductions of FA observed in aging are thought to reflect general degeneration of myelinated fibers in white matter (Salat 2011; Bennett and Madden 2014), although the specific pathology of this process is under debate. Greater white matter tissue anisotropy may reflect more intact axonal quality (e.g., degree of myelination, fiber thickness) and/or quantity (e.g., number of fibers), all of which affect the efficiency of neuronal signal transmission between cortical areas connected by white matter. In line with this interpretation, we hypothesized that flexible modulation of neural activity under greater cognitive demands would be affected by age-related declines in microstructural integrity, and that this association should influence cognitive performance.

Our results supported this hypothesis, with both FA and modulation of BOLD activity together accounting for age-related variability in cognitive performance. This provides important evidence for the assumption that declines in cognitive ability with age are partly a result of both structural degradation and functional dysregulation. These findings further highlight the importance of the maintenance of both white matter connectivity and flexible neural modulation in support of cognitive performance with aging. Beyond the combined effects of FA and positive and negative functional modulation, unique influences of structure on the direction of functional modulation were observed. FA explained age differences in task performance indirectly through its effects on upregulating functional responses to increased task demands. Specifically, greater age was associated with reductions in FA, and lower FA was related to decreases in the degree of positive BOLD modulation, which predicted poorer cognitive performance (both during *n*-back and on tests of executive function). Our results provide multivariate evidence from a wide age range of individuals that age-related degradation of white matter tracts reduces the ability to upregulate cortical activity in regions supporting response to cognitive challenge, and consequently, results in poorer cognitive performance. Degraded white matter likely results in disrupted organization/propagation of neural signaling, contributing to inefficient neural responses to changes in task demands. This, in turn negatively impacts working memory task performance and cognitive ability more generally. Associations between white matter FA and level of neural activity in both task-related and default-mode regions have been previously demonstrated in aging (Madden et al. 2007; de Chastelaine et al. 2011; Brown et al. 2015, 2018; Daselaar et al. 2015; Zhu et al. 2015). Additionally, in a small group of older adults, three year change in FA of the body of the corpus callosum was associated with altered activation in a prefrontal region associated with task performance (Hakun et al. 2015). Our results support this work, showing that age-related degradation of white matter connectivity across the full adult lifespan affects the magnitude of neural activity. Moreover, we extend existing reports by demonstrating that this type of structure-function relationship influences the level of both working memory task performance and general executive functioning in healthy aging.

Interestingly, negative modulation was found to account for variance in cognitive performance due to aging, yet independent of a significant relationship with FA. Given previous work linking reductions in FA to weakened deactivation in DMN regions (Brown et al. 2015, 2018), it is not clear why in our sample age-related alterations in FA were related to the degree of positive, but not negative modulation. One reason may be the fact that activation in our study represented parametric modulation associated with differing levels of task difficulty, as opposed to explicit identification of the magnitude of task-negative (de)activation. Further work is needed to clarify the effects of declining brain structure on regulation of functional responses to cognitive challenge across various tasks.

Overall, our results support a cortical disconnection theory of cognitive aging which posits that age-related declines in cognition occur, in part, as a result of breakdowns in both structural components of the brain and in functional responses to cognitive challenge (O’Sullivan et al. 2001; Bartzokis 2004). This theory assumes that cognitive decline occurs in healthy aging as a result of “disconnected” structural brain systems/networks, constraining function of gray matter integrated within these networks. Here, we demonstrate that age-related white matter degradation and dysregulation of functional systems are jointly associated with poorer cognitive performance, and more specifically that decrements to structural connection reduce upregulation of functional regions to meet task demands. Importantly, the fact that this theorized structure-function association also explained variation in executive function differences across age supports and strengthens the idea that disruption of brain systems represents a more generalized picture of age-related cognitive decline. While there are likely alternative facets of brain structure that alter BOLD modulation, our results further highlight the importance of maintaining structural brain integrity to support dynamic range of functional modulation and cognitive functioning in older age.

The present findings should be interpreted in the context of study limitations. While SEM provides a structure with which to test theoretical frameworks of cognitive aging, it is still not possible to make definitive causal links between age, brain, and cognitive variables, especially in a cross-sectional design. Longitudinal data are required to draw conclusions about how the process of aging influences individual brain structure-function relationships, and how that ultimately affects cognitive processes (Lindenberger et al. 2011). It is also important to note that DTI is an indirect proxy of white matter microstructure, and thus, is not a direct measure of axonal structure or integrity. However, the use of tractography methods provides more precise information on white matter organization beyond that obtained in previous studies using region of interest approaches. Lastly, we cannot speak to specificity in relationships between particular white matter tracts and regions of gray matter activation; however, the use of latent variables to quantify white matter integrity, functional modulation, and task performance is a strength of our approach. Through this, we were able to reduce the chance that identified structure-function relationships were due to specificity in localized tracts or ROIs, allowing for a more generalized perspective of structure-function effects on cognitive aging. Lastly, a caveat of any SEM analysis, the tested hypothesis is just one possible theoretical account of the relationship of these *a priori* selected variables of study. There remains variance to be accounted for that could be specified in other models and that could include a myriad of other factors salient to the aging process (e.g., gray matter volume/thickness/surface area, white matter lesions, beta-amyloid deposition, iron accumulation, vascular health risk, and so forth). Future work, with even larger samples, should aim to deduce the influential relationships among these additional biological variables on structure-function interactions that contribute to cognitive decline.

## Conclusion

A wealth of previous research demonstrates that typical aging is associated with disrupted neural systems. Declining structural connectivity and impaired neuronal communication in aging are assumed to contribute to altered functional activation patterns, which ultimately result in reduced cognitive functioning. To our knowledge, this is the first study to simultaneously establish structure-function-cognition associations across the lifespan. Here, we demonstrate that aging affects both microstructural components of white matter and functional modulation of neural activity, and importantly, that age-related reductions in these structural and functional components have negative cognitive consequences. More specifically, our results suggest that reductions in the ability to upregulate functional activation in response to cognitive challenge negatively affects cognition, partly as a result of age-related impairments in neuronal signaling from disrupted structural connectivity. Therefore, our study highlights the importance of structural properties in the regulation of neural activity and cognitive function across the lifespan, and lends novel support to theories of cortical “disconnection” as a plausible mechanism of age-related cognitive decline.

## References

Bartzokis G. 2004. Age-related myelin breakdown: a developmental model of cognitive decline and Alzheimer’s disease. Neurobiol Aging. 25:5–18.

Bennett IJ, Madden DJ. 2014. Disconnected aging: Cerebral white matter integrity and age-related differences in cognition. Neuroscience. 276:187–205.

Bennett IJ, Rypma B. 2013. Advances in functional neuroanatomy: A review of combined DTI and fMRI studies in healthy younger and older adults. Neurosci Biobehav Rev. 37:1201–1210.

Borghesani PR, Madhyastha TM, Aylward EH, Reiter MA, Swarny BR, Warner Schaie K, Willis SL. 2013. The association between higher order abilities, processing speed, and age are variably mediated by white matter integrity during typical aging. Neuropsychologia. 51:1435–1444.

Brett M, Anton JL, Valabregue R, Poline JB. 2002. Imaging techniques. Neuroimage. 16:37–258.

Brickman AM, Meier IB, Korgaonkar MS, Provenzano FA, Grieve SM, Siedlecki KL, Wasserman BT, Williams LM, Zimmerman ME. 2012. Testing the white matter retrogenesis hypothesis of cognitive aging. Neurobiol Aging. 33:1699–1715.

Brown CA, Hakun JG, Zhu Z, Johnson NF, Gold BT. 2015. White matter microstructure contributes to age-related declines in task-induced deactivation of the default mode network. Front Aging Neurosci. 7:194.

Brown CA, Jiang Y, Smith CD, Gold BT. 2018. Age and Alzheimer’s pathology disrupt default mode network functioning via alterations in white matter microstructure but not hyperintensities. Cortex. 104:58–74.

Cabeza R, Nyberg L. 2000. Imaging Cognition II: An Empirical Review of 275 PET and fMRI Studies. J Cogn Neurosci. 12:1–47.

Cappell KA, Gmeindl L, Reuter-Lorenz PA. 2010. Age differences in prefontal recruitment during verbal working memory maintenance depend on memory load. Cortex. 46:462–473.

Daselaar SM, Iyengar V, Davis SW, Eklund K, Hayes SM, Cabeza RE. 2015. Less Wiring, More Firing: Low-Performing Older Adults Compensate for Impaired White Matter with Greater Neural Activity. Cereb Cortex. 25:983–990.

de Chastelaine M, Wang TH, Minton B, Muftuler LT, Rugg MD. 2011. The Effects of Age, Memory Performance, and Callosal Integrity on the Neural Correlates of Successful Associative Encoding. Cereb Cortex. 21:2166–2176.

Delis DC, Kaplan E, Kramer JH. 2001. Delis-Kaplan executive function system. San Antonio, TX Psychol Corp.

Desikan RS, Ségonne F, Fischl B, Quinn BT, Dickerson BC, Blacker D, Buckner RL, Dale AM, Maguire RP, Hyman BT, Albert MS, Killiany RJ. 2006. An automated labeling system for subdividing the human cerebral cortex on MRI scans into gyral based regions of interest. Neuroimage. 31:968–980.

Fjell AM, Walhovd KB. 2010. Structural Brain Changes in Aging: Courses, Causes and Cognitive Consequences. Rev Neurosci. 21:187–222.

Folstein MF, Folstein SE, McHugh PR. 1975. “Mini-mental state”. A practical method for grading the cognitive state of patients for the clinician. J Psychiatr Res. 12:189– 198.

Garrett DD, Kovacevic N, McIntosh AR, Grady CL. 2013. The Modulation of BOLD Variability between Cognitive States Varies by Age and Processing Speed. Cereb Cortex. 23:684–693.

Gold BT, Powell DK, Xuan L, Jicha GA, Smith CD. 2010. Age-related slowing of task switching is associated with decreased integrity of frontoparietal white matter. Neurobiol Aging. 31:512–522.

Gusnard DA, Raichle ME. 2001. Searching for a baseline: Functional imaging and the resting human brain. Nat Rev Neurosci. 2:685–694.

Hakun JG, Johnson NF. 2017. Dynamic range of frontoparietal functional modulation is associated with working memory capacity limitations in older adults. Brain Cogn. 118:128–136.

Hakun JG, Zhu Z, Brown CA, Johnson NF, Gold BT. 2015. Longitudinal alterations to brain function, structure, and cognitive performance in healthy older adults: A fMRI-DTI study. Neuropsychologia.

Hofer S, Frahm J. 2006. Topography of the human corpus callosum revisited— Comprehensive fiber tractography using diffusion tensor magnetic resonance imaging. Neuroimage. 32:989–994.

Hu L, Bentler PM. 1999. Cutoff criteria for fit indexes in covariance structure analysis: Conventional criteria versus new alternatives. Struct Equ Model A Multidiscip J. 6:1–55.

Hua K, Zhang J, Wakana S, Jiang H, Li X, Reich DS, Calabresi PA, Pekar JJ, van Zijl PCM, Mori S. 2008. Tract probability maps in stereotaxic spaces: Analyses of white matter anatomy and tract-specific quantification. Neuroimage. 39:336–347.

Kennedy KM, Boylan MA, Rieck JR, Foster CM, Rodrigue KM. 2017. Dynamic range in BOLD modulation: lifespan aging trajectories and association with performance. Neurobiol Aging. 60:153–163.

Kennedy KM, Raz N. 2009. Aging white matter and cognition: Differential effects of regional variations in diffusion properties on memory, executive functions, and speed. Neuropsychologia. 47:916–927.

Kennedy KM, Rodrigue KM, Bischof GN, Hebrank AC, Reuter-Lorenz PA, Park DC. 2015. Age trajectories of functional activation under conditions of low and high processing demands: An adult lifespan fMRI study of the aging brain. Neuroimage. 104:21–34.

Kline RB. 2011. Principles and Practice of Structural Equation Modeling, Guilford Publications.

Leemans A, Jones DK. 2009. The B-matrix must be rotated when correcting for subject motion in DTI data. Magn Reson Med. 61:1336–1349.

Lindenberger U, von Oertzen T, Ghisletta P, Hertzog C. 2011. Cross-sectional age variance extraction: What’s change got to do with it? Psychol Aging. 26:34–47.

MacCallum RC, Browne MW, Sugawara HM. 1996. Power analysis and determination of sample size for covariance structure modeling. Psychol Methods. 1:130–149.

Madden DJ, Bennett IJ, Song AW. 2009. Cerebral White Matter Integrity and Cognitive Aging: Contributions from Diffusion Tensor Imaging. Neuropsychol Rev. 19:415– 435.

Madden DJ, Spaniol J, Costello MC, Bucur B, White LE, Cabeza R, Davis SW, Dennis NA, Provenzale JM, Huettel SA. 2009. Cerebral White Matter Integrity Mediates Adult Age Differences in Cognitive Performance. J Cogn Neurosci. 21:289–302.

Madden DJ, Spaniol J, Whiting WL, Bucur B, Provenzale JM, Cabeza R, White LE, Huettel SA. 2007. Adult age differences in the functional neuroanatomy of visual attention: A combined fMRI and DTI study. Neurobiol Aging. 28:459–476.

Mazaika P, Whitfield-Gabrieli S, Reiss A. 2007. Artifact repair for fMRI data from high motion clinical subjects. Poster Session Presented at Human Brain Mapping. Chicago, IL.

Mori S, Oishi K, Jiang H, Jiang L, Li X, Akhter K, Hua K, Faria A V., Mahmood A, Woods R, Toga AW, Pike GB, Neto PR, Evans A, Zhang J, Huang H, Miller MI, van Zijl P, Mazziotta J. 2008. Stereotaxic white matter atlas based on diffusion tensor imaging in an ICBM template. Neuroimage. 40:570–582.

Muthen LK, Muthen BO. 2017. Mplus User’s Guide. Seventh Edition.

Nagel IE, Preuschhof C, Li S-C, Nyberg L, Bäckman L, Lindenberger U, Heekeren HR. 2011. Load Modulation of BOLD Response and Connectivity Predicts Working Memory Performance in Younger and Older Adults. J Cogn Neurosci. 23:2030– 2045.

O’Sullivan M, Jones DK, Summers PE, Morris RG, Williams SCR, Markus HS. 2001. Evidence for cortical “disconnection” as a mechanism of age-related cognitive decline. Neurology. 57:632–638.

Oguz I, Farzinfar M, Matsui J, Budin F, Liu Z, Gerig G, Johnson HJ, Styner M. 2014. DTIPrep: quality control of diffusion-weighted images. Front Neuroinform. 8:4.

Owen AM, McMillan KM, Laird AR, Bullmore E. 2005. N-back working memory paradigm: A meta-analysis of normative functional neuroimaging studies. Hum Brain Mapp. 25:46–59.

Park D. 2010. Age differences in default mode activity on easy and difficult spatial judgment tasks. Front Hum Neurosci. 3:75.

Peirce JW. 2008. Generating stimuli for neuroscience using PsychoPy. Front Neuroinform. 2:10.

Persson J, Lustig C, Nelson JK, Reuter-Lorenz PA. 2007. Age Differences in Deactivation: A Link to Cognitive Control? J Cogn Neurosci. 19:1021–1032.

Persson J, Nyberg L, Lind J, Larsson A, Nilsson L-G, Ingvar M, Buckner RL. 2006. Structure–Function Correlates of Cognitive Decline in Aging. Cereb Cortex. 16:907–915.

Radloff LS. 1977. The CES-D Scale. Appl Psychol Meas. 1:385–401.

Raichle ME, MacLeod AM, Snyder AZ, Powers WJ, Gusnard DA, Shulman GL. 2001. A default mode of brain function. Proc Natl Acad Sci. 98:676–682.

Raz N, Kennedy KM. 2009. A Systems Approach to the Aging Brain: Neuroanatomic Changes, Their Modifiers, and Cognitive Correlates. In: Imaging the Aging Brain. Oxford University Press. p. 43–70.

Rieck JR, Rodrigue KM, Boylan MA, Kennedy KM. 2017. Age-related reduction of BOLD modulation to cognitive difficulty predicts poorer task accuracy and poorer fluid reasoning ability. Neuroimage. 147:262–271.

Salat DH. 2004. Thinning of the Cerebral Cortex in Aging. Cereb Cortex. 14:721–730.

Salat DH. 2011. The Declining Infrastructure of the Aging Brain. Brain Connect. 1:279– 293.

Samanez-Larkin GR, Levens SM, Perry LM, Dougherty RF, Knutson B. 2012. Frontostriatal White Matter Integrity Mediates Adult Age Differences in Probabilistic Reward Learning. J Neurosci. 32:5333–5337.

Sambataro F, Murty VP, Callicott JH, Tan H-Y, Das S, Weinberger DR, Mattay VS. 2010. Age-related alterations in default mode network: Impact on working memory performance. Neurobiol Aging. 31:839–852.

Schneider-Garces NJ, Gordon BA, Brumback-Peltz CR, Shin E, Lee Y, Sutton BP, Maclin EL, Gratton G, Fabiani M. 2010. Span, CRUNCH, and Beyond: Working Memory Capacity and the Aging Brain. J Cogn Neurosci. 22:655–669.

Sullivan E V., Pfefferbaum A. 2006. Diffusion tensor imaging and aging. Neurosci Biobehav Rev. 30:749–761.

Turner GR, Spreng RN. 2015. Prefrontal Engagement and Reduced Default Network Suppression Co-occur and Are Dynamically Coupled in Older Adults: The Default– Executive Coupling Hypothesis of Aging. J Cogn Neurosci. 27:2462–2476.

Witelson SF. 1989. Hand and sex differences in the isthmus and genu of the human corpus callosum. A postmortem morphological study. Brain. 112:799–835.

Yeh F-C, Verstynen TD, Wang Y, Fernández-Miranda JC, Tseng W-YI. 2013. Deterministic Diffusion Fiber Tracking Improved by Quantitative Anisotropy. PLoS One. 8:e80713.

Zhu Z, Johnson NF, Kim C, Gold BT. 2015. Reduced Frontal Cortex Efficiency is Associated with Lower White Matter Integrity in Aging. Cereb Cortex. 25:138–146.

